# Cln3 can work independently of Whi5 on the cell size for Start in yeast

**DOI:** 10.1101/2025.10.24.684447

**Authors:** Amrita Shah, Sangeet Honey, Richard Sejour, Chenjun He, Bruce Futcher

## Abstract

In yeast, cells commit to division at “Start” in G1 phase. The critical cell size for Start is regulated in part by the size-control genes *CLN3* and *WHI5*. Cln3 is a G1 cyclin that activates Start by activating the Cdc28 cyclin dependent kinase. Null *cln3* mutants have large cells, because they lack an activator, while hyperactive mutants have small cells. Whi5 inhibits Start. Null *whi5* mutants have small cells, while hyperactive mutants have large cells. Previous studies suggested that *CLN3* and *WHI5* are in a linear, dependent pathway, in which Cln3 acts inhibits Whi5, possibly by phosphorylation. We compared isogenic WT, *cln3, whi5*, and *cln3 whi5* mutants, and find that *whi5* is not epistatic, or not fully epistatic, to *CLN3*—that is, *CLN3* can work independently of *WHI5*. The *cln3* and *whi5* deletions have off-setting phenotypes, such that the double mutant is nearly wild-type in cell size, and has only the mildest phenotype, with well-controlled cell size, despite entirely lacking two major cell size regulators. Growth rates were also measured. Some mutants were larger and some mutants smaller than wild-type, but the wild-type had the fastest doubling time, consistent with the idea that wild-type size is optimal for growth rate.

## Introduction

Growing, dividing cells maintain an equilibrium distribution of cell sizes from generation to generation, and this implies a mechanism to co-ordinate division with growth. In general, it is believed that cells must grow to some sufficient size, and this then allows division (Johnston, Pringle and Hartwell 1977). The mechanism of this linkage is still unclear, but in 1975 Nurse proposed that mutants that altered the equilibrium cell size could report on this linkage (Nurse 1975). The *S. pombe cdc9-50* (a.k.a. *wee1*) mutant found by Nurse, and many other cell size mutants subsequently found by other workers, turned out to affect central components of the cell cycle machinery. For instance, the *S. pombe* mutant *wee1* is in a tyrosine kinase that phosphorylates and regulates Cdc2, the Cyclin Dependent Kinase (CDK) that drives the cell cycle, while the *wee2* mutant is a mutation in the CDK itself (Nurse and Thuriaux 1980). Both these mutants have unusually small cells. In *S. cerevisiae*, the dominant, hyperactive mutant *WHI1-1* (now known as *CLN3-1*) defined the first G1 cyclin, which helps activate Cdc28, the cell division CDK, and the *cln3* null mutant has large cells (Sudbery, Goodey and Carter 1980; Nash *et al*. 1988), as expected for an activator of cell cycle entry.

In *S. cerevisiae*, two of the best known cell size control genes are *CLN3* (Sudbery, Goodey and Carter 1980) and *WHI5* (Jorgensen *et al*. 2002), which work in opposite directions. A *cln3* mutant has large cells (Nash *et al*. 1988), while a *whi5* mutant has small cells (Jorgensen *et al*. 2002). Jorgensen et al. crossed these mutants to make a *cln3 whi5* double mutant; no data were shown, but the genes were described as additive, not epistatic, though the definition of epistasis in that paper was quite demanding (Jorgensen *et al*. 2002). Later, Costanzo et al. and de Bruin et al. did molecular and biochemical characterizations of *WHI5* and its encoded protein (Costanzo *et al*. 2004; De Bruin *et al*. 2004). Each paper showed one cell size plot of the four possible genotypes (*CLN3 WHI5, cln3 WHI5, CLN3 whi5*, and *cln3 whi5*). The plots in the two papers were somewhat at odds with each other; Costanzo et al. described *whi5* as partly epistatic to *CLN3*, whereas de Bruin et al found that the *cln3 whi5* double mutant cells had the same size as *whi5* cells (i.e., both small), and accordingly described *whi5* as epistatic to *CLN3*. Since *CLN3* is an activator of the cell cycle (the cell cycle commitment point is called “Start” in yeast), and *WHI5* is an inhibitor, it was natural to interpret this epistasis as *CLN3* being an inhibitor of *WHI5*, and *WHI5* as an inhibitor of Start, the commitment to the cell cycle. Accordingly, some ensuing literature has used simplified models where the sole role of *CLN3* is to inhibit *WHI5*, and this can be seen in the introduction to many cell cycle papers, and in reviews.

More recently, Wang et al. found that over-expression of *CLN3* from a *GAL* promoter caused smaller cell size even in a *whi5* deletion mutant (Wang *et al*. 2009), inconsistent with the idea that *CLN3* works solely via *WHI5*. This and other related results in our laboratory, prompted us to re-examine the relationship between *CLN3* and *WHI5*. We made isogenic *cln3* and *whi5* mutants, crossed them, and dissected tetrads, obtaining many tetratypes (i.e., *CLN3 WHI5, cln3 WHI5, CLN3 whi5*, and *cln3 whi5* tetrads). We measured cell sizes in many isogenic tetrads, in two genetic backgrounds. Ultimately, our results agree with those of Costanzo et al. (Costanzo *et al*. 2004): the two genes can work independently, with *whi5* only slightly epistatic to *CLN3*, implying that *CLN3* has additional targets.

*CLN3* and *WHI5* are sometimes discussed as the two major cell size control genes in yeast (Litsios *et al*. 2022; Schmoller *et al*. 2022). And yet, we show below that the *cln3 whi5* double null mutant has only a very mild phenotype: it is nearly wild-type in size; its morphology is nearly wild-type; the co-efficient of variation for cell size is the same as wild-type; and its growth rate is nearly wild-type. Yeast apparently do very well even when lacking both of these size control genes.

Finally, it is still an open question why different kinds of cells have the particular sizes they do. Since we had many sets of isogenic strains with the four possible genotypes, and each genotype had a different size, both smaller and larger than wild-type, we measured growth rates in many isogenic strains, finding that the wild-type has the fastest growth.

## Methods

### Strains

Most experiments were done in the GZ background (Zhao *et al*. 2016). GZ240 (MATa) and GZ241 (MATalpha) are quasi-wildtype strains. They come from a cross between DBY12032 (D. Botstein) and UCC8378 (Dimitrov *et al*. 2009), both in the S288C background. Compared to S288C, they differ mainly in carrying the fully wild-type *HAP1* allele, and the mitochondria-stabilizing alleles of *SAL1, CAT5, MIP1, MKT1* found in UCC8376. These are robustly-growing strains with well-regulated respiration; they grow somewhat faster than the common lab strains BY4741 and BY4742.

GZ240 and GZ241 are prototrophic. Derivatives deleted for *ura3* and *leu2* (GZ238 MATa, GZ239 MATalpha) were created for genetic analysis (Zhao *et al*. 2016). The AS series of strains used below, carrying alleles of *cln3* and *whi5*, were derived directly from GZ238 and GZ239, and are *leu2 ura3*.

We also made prototrophic versions of these strains by crossing to GZ240 and GZ241, yielding prototrophic versions of the *CLN3 WHI5, cln3 WHI5, CLN3 whi5*, and *cln3 whi5* strains. Prototrophs were used in one of the three growth rate experiments, E1 (Table 5).

In addition, we did experiments in the BY4741/BY4742 background of the yeast deletion set. In these experiments, the *whi5::G418* allele was derived from the deletion set. The *cln3::G418* allele was derived from the BY4742 (MATalpha) strain of the deletion set. We believe the *cln3* allele in the BY4741 (MATa) strain of the deletion set may not be correct. Mutant combinations were obtained by crossing BY4741/BY4742 strains from the deletion set, and picking tetrads.

### Deleting *CLN3* and *WHI5* in the GZ background

Synthetic DNA fragments were made at Twist Bioscience to delete *CLN3* and *WHI5*. These fragments contain about 200 bp of 5’ and 3’ flanking homology to *CLN3* or *WHI5*, respectively; inside these regions of flanking homology they contain a KanMX expression cassette, which is flanked by 130 bp direct repeats of a sequence originating from the *hisG* gene of *E. coli*. The intention of this arrangement is that integration of the whole cassette is targeted by the 200 bp of flanking homology, with selection for resistance to G418. But subsequently, after a Cas9-induced cut within the KanMX sequence, recombination between the 130 bp *hisG* repeats removes the KanMX expression cassette.

GZ238 and GZ239 were transformed with the *CLN3* or *WHI5* disrupting fragment, and colonies resistant to G418 were selected. These were assayed by PCR and by cell size phenotype (using a Coulter Channelizer) to confirm that the transformants were *cln3* or *whi5* mutants. Several such mutants were then transformed with a plasmid carrying *LEU2* and Cas9, and a guide RNA targeting the KanMX cassette at the sequence TTACTCACCACTGCGATCCC. After Cas9-mediated cutting, many transformed cells healed the cut by recombination between the two 130 bp direct repeats of *hisG*. These were selected on -leu plates, selecting for the *LEU2* Cas9 plasmid. Cells that did not heal the cut did not survive. Transformants were assayed by PCR for the expected presence of a single 130 bp *hisG* insertion replacing the open reading frame of *CLN3* or *WHI5*. Several such transformants were chosen, and their size phenotypes were assayed using a Coulter Channelizer. Thus these strains had their *CLN3* or *WHI5* genes replaced by a 130 bp sequence from *hisG*, and no longer contained KanMX sequences. These deletion constructs are independent of and different from the deletions in the BY4741/4742 background, arguing that the exact nature of the null allele does not significantly affect the phenotype.

Strains with a *cln3::hisG* deletion were crossed to strains with a *whi5::hisG* deletion. Diploids were chosen by microdissection. Diploids were sporulated, and tetrads were dissected. Sporulation was high and spore viability was high, and rare inviability did not seem associated with any particular genotype. Inspection of the tetrad dissection plates showed that all growing spore clone colonies were very similar in size, suggesting that none of the allele combinations caused any significant growth defect. 2:2 segregation of the *cln3::hisG* allele and the *whi5::hisG* allele was confirmed by PCR. Also segregation of cell size was assayed using a Coulter Channelizer. The predicted cell sizes were seen for the WT, *cln3*, and *whi5* spore clones, while the cell sizes of the *cln3 whi5* double mutants were the main research question.

In the BY background, the *cln3:KanMX* and *whi5:KanMX* alleles were taken from the yeast deletion set. The *cln3::KanMX* allele came from the *MATalpha* strain of the yeast deletion set; we believe there is something wrong with the *cln3::KanMX* allele in the *MATa* strain.

### Media

For size and growth rate measurements, cells were grown in YEPD (Yeast Extract Peptone Dextrose). This was 1% yeast extract (Gibco/Thermo Fisher), 2% Gibco Soy Peptone 100 with supplements (see below), 2% D-glucose. For YEPD, Difco Bacto Peptone has traditionally been used. This is a digest of bovine and porcine proteins. Instead, we use a peptone from soy bean proteins. However, this is somewhat low in certain amino acids compared to an animal peptone. Therefore, we supplement each 500 g container of Soy Peptone 100 with 2.5 g each of leucine, lysine, histidine, methionine, cysteine, and tryptophan, and 0.25 g of adenine. Typical yeast strains (BY background, GZ background with or without auxotrophies) grow somewhat faster in this YEPD with supplemented Soy Peptone than in YEPD with regular Bacto Peptone.

### Instrumentation

For measuring cell size, and for counting cell numbers, in the early part of this work, we used a Beckman-Coulter Z2 Coulter Counter (“Channelizer”). In the later part of the work, we used a Beckman-Coulter Multisizer 4e. Channelizer data were used in Fig. 2B, and in Table 3; all other data were generated with the Multisizer 4e. Both instruments used a 50 micron aperture, and both were calibrated with Beckman-Coulter calibration beads. Cell volumes are reported in femtolitres (fL).

## Results

### Cell Size Distributions

**Fig. 1** shows a typical overlay of the cell size distributions for the four genotypes from three tetratype tetrads in the GZ background. All four genotypes have distinguishable cell size distributions, and it appears that *cln3* and *whi5* are not epistatic to each other. That is, the *cln3* deletion has an effect even in a *whi5* strain; and a *whi5* deletion has an effect even in a *cln3* strain. The *cln3 whi5* double mutant is very similar to wild-type, even though it is completely lacking two major cell size regulators.

**Figure 1.**
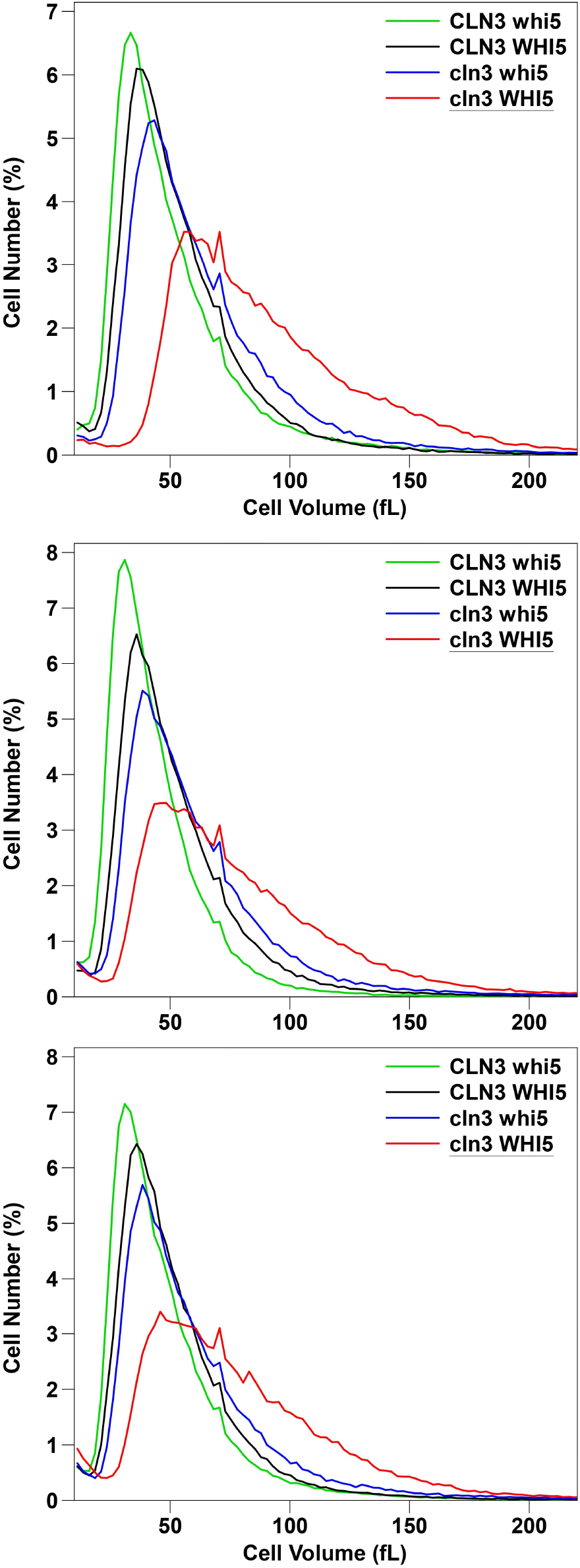
From top to bottom, Fig. 1A, 1B, and 1C, showing AS tetrads 9, 10, and 12, respectively. Identities, genotypes, and mean, median, and mode cell sizes of each spore clone are shown in Table S1. Cells were grown to early log phase (∼ 2 x 10^7 cells/ml) and cell sizes in fL were measured using a Beckman-Coulter Multisizer 4e with a 50 micron aperture.

Figure 2 summarizes quantitative mean cell size data for the four genotypes in both the GZ background and the BY background. Full data, including individual measurements of mean, median and mode, and error data, are in supplementary results (Table S1, S2). Fig. 2 confirms the impression of Fig. 1 that *whi5 cln3* double mutants are larger than *whi5* single mutants, and thus that *whi5* is not epistatic, or not fully epistatic, to *cln3*.

**Figure 2.**
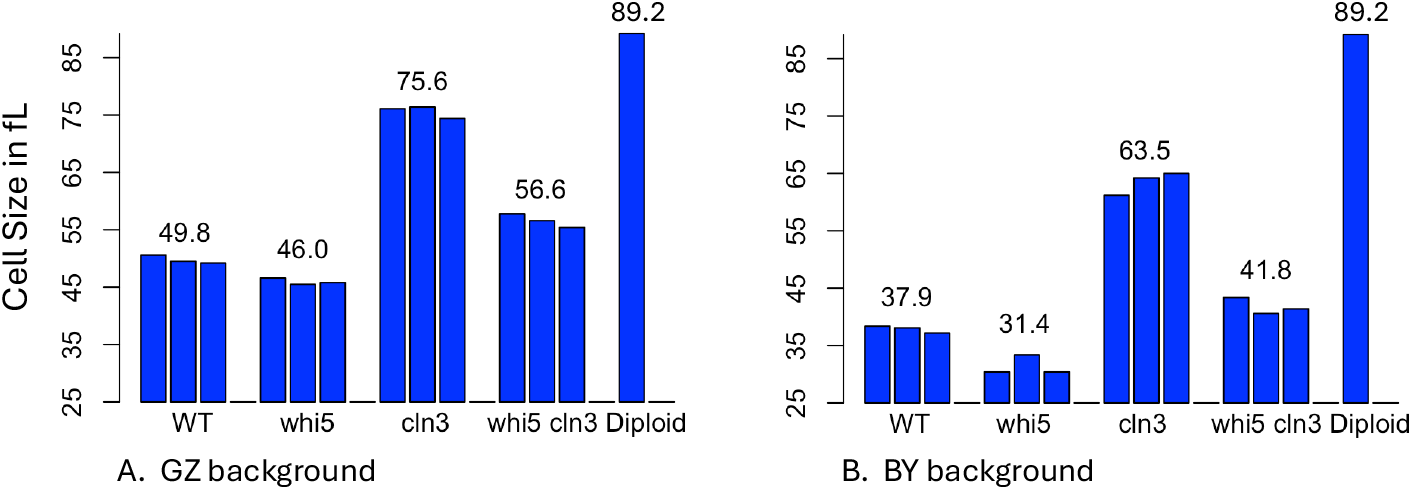
Mean Cell Sizes

Figure 2 Legend. Fig. 2A. Cells were grown to early log phase (∼2 x 10^7 cells/ml) in YEPD. Cell sizes were measured with a Beckman-Coulter Multisizer with a 50 micron aperture, and mean cell sizes are shown. Each bar represents the mean of six technical replicates of a single strain of the stated genotype. Thus, the three wild-type bars represent a total of 18 measurements of three different but isogenic wild-type strains. The strains come from the AS9, AS10, and AS12 tetrads. The diploid was in the GZ background, and was a cln3/+ whi5/+ double heterozygote. Standard errors are shown numerically in Table S1. Fig. 2B. As Fig. 2A, but with strains from the BY background, measured with a Beckman-Coulter Z2 Coulter Counter with a 50 micron aperture. In this case, only 1 technical replicate was done. Numerical data is shown in Table S2. In 2A and 2B, the two diploid bars shown are not independent measurements, but are the same data added to both graphs for comparison.

**Table 1** summarizes size results for each genotype. In both genetic backgrounds, the *cln3* mutant has the biggest cells, followed by the double mutant, followed by the wild-type, followed by the *whi5* mutant. Each mutant combination has a statistically-different size from each of the other combinations, with very small p-values. Notably, the *cln3 whi5* double mutant has a different size from both the *cln3* single mutant and the *whi5* single mutant. This is inconsistent with models in which Cln3 and Whi5 act in a single, linear pathway, in which case one gene would be epistatic to the other, and the double mutant would be expected to have the same size as one (or perhaps both) of the single mutants.

**Table 1.** Mean cell sizes with (standard error)

| Back-ground | N | Genotype | Mean Size, fL (SE) | p-value vs. |  |  |
| --- | --- | --- | --- | --- | --- | --- |
|  |  |  |  | *CLN3 WHI5* | *CLN3 whi5* | *cln3 WHI5* |
| GZ238/<br>GZ239 | 18 | *CLN3 WHI5* | 49.7 (0.20) |  |  |  |
|  | 18 | *CLN3 whi5* | 46.0 (0.18) | 8e-16 |  |  |
|  | 18 | *cln3 WHI5* | 75.6 (0.29) | 3e-39 | 1e-41 |  |
|  | 18 | *cln3 whi5* | 56.6 (0.32) | 3e-19 | 9e-26 | 9e-32 |
|  | 8 | *Diploid* | 89.2 (0.38) |  |  |  |
| BY4741/<br>BY4742 | 3 | *CLN3 WHI5* | 37.88 |  |  |  |
|  | 3 | *CLN3 whi5* | 31.41 | 4e-3 |  |  |
|  | 3 | *cln3 WHI5* | 63.45 | 3e-5 | 3e-5 |  |
|  | 3 | *cln3 whi5* | 41.78 | 1e-2 | 1e-3 | 1e-4 |

**Table 2** shows the coefficient of variation (CV) for the cell size distribution for each genotype, assuming (incorrectly) that cell size has a normal distribution. (Alternative methods for comparing size distributions are discussed by (Chen *et al*. 2020)) If one were to statistically compare these CVs based on a normal distribution, all pairwise comparisons would be significant. However, since the size distributions are not normal, this is not an appropriate test. Interestingly, the CV for the wild-type and for the double mutant are very similar, despite the fact that the double mutant lacks two major cell size controllers.

**Table 2.**

| Back-ground | N | Genotype | CV (size, fL) |
| --- | --- | --- | --- |
| GZ238/<br>GZ239 | 18 | *CLN3 WHI5* | 0.47 |
|  | 18 | *CLN3 whi5* | 0.54 |
|  | 18 | *cln3 WHI5* | 0.44 |
|  | 18 | *cln3 whi5* | 0.49 |
|  | 8 | *Diploid* | 0.37 |

**Table 3.** Cells were grown to early log phase and sized with a Coulter Counter Z2. The proportion of cells larger than the mean plus two standard deviations is shown.

| Genotype | % > (mean + 2SD) |
| --- | --- |
| *CLN3 whi5* | 4.93% |
| *CLN3 WHI5* | 4.31% |
| *cln3 whi5* | 5.05% |
| *cln3 WHI5* | 4.73% |

We can compare the observed size of the double mutant to the size expected if *CLN3* and *WHI5* work independently. The expected size of the double mutant (based on independence) can be calculated as (*whi5* proportion of WT size) x (*cln3* proportion of WT size) x (WT size) = ?? fL. For the GZ background, this expected size of the double mutant is 69.9 fL, vs 56.6 observed (observed is 81% of expected), and for the BY background, the expected size is 52.6 fL, vs 41.8 observed (observed is 79% of expected). Thus the double mutants are about 20% smaller than expected on the basis of independence, arguing that the *whi5* mutation has the larger effect (i.e., that it is partly but not completely epistatic to *CLN3*, as argued by Costanzo et al. (Costanzo *et al*. 2004)). (A similar result is obtained in an additive rather than a multiplicative model.) This result could be interpreted to mean that Cln3 does work, in part, via Whi5, but works via other pathways as well. Although it may be valuable to make this comparison, the argument for “partial epistasis” is not rigorous.

### Qualitative effects

The *cln3* and *whi5* mutants have off-setting phenotypes, and indeed the *cln3 whi5* mutant is nearly the same size as wild-type (**Fig. 1, Fig. 2, Table 1**). Under the microscope, the *cln3 whi5* double mutant is almost indistinguishable from wild-type with no aberrant morphology (**Fig 3**); the overall cell size distribution is very similar to wild-type; and the doubling time is also very close to wild-type (see below). It seems that yeast cells do a very good job of controlling cell size and cell division even in the complete absence of both of these important cell size controllers. Presumably there are other redundant controls.

**Figure 3.**
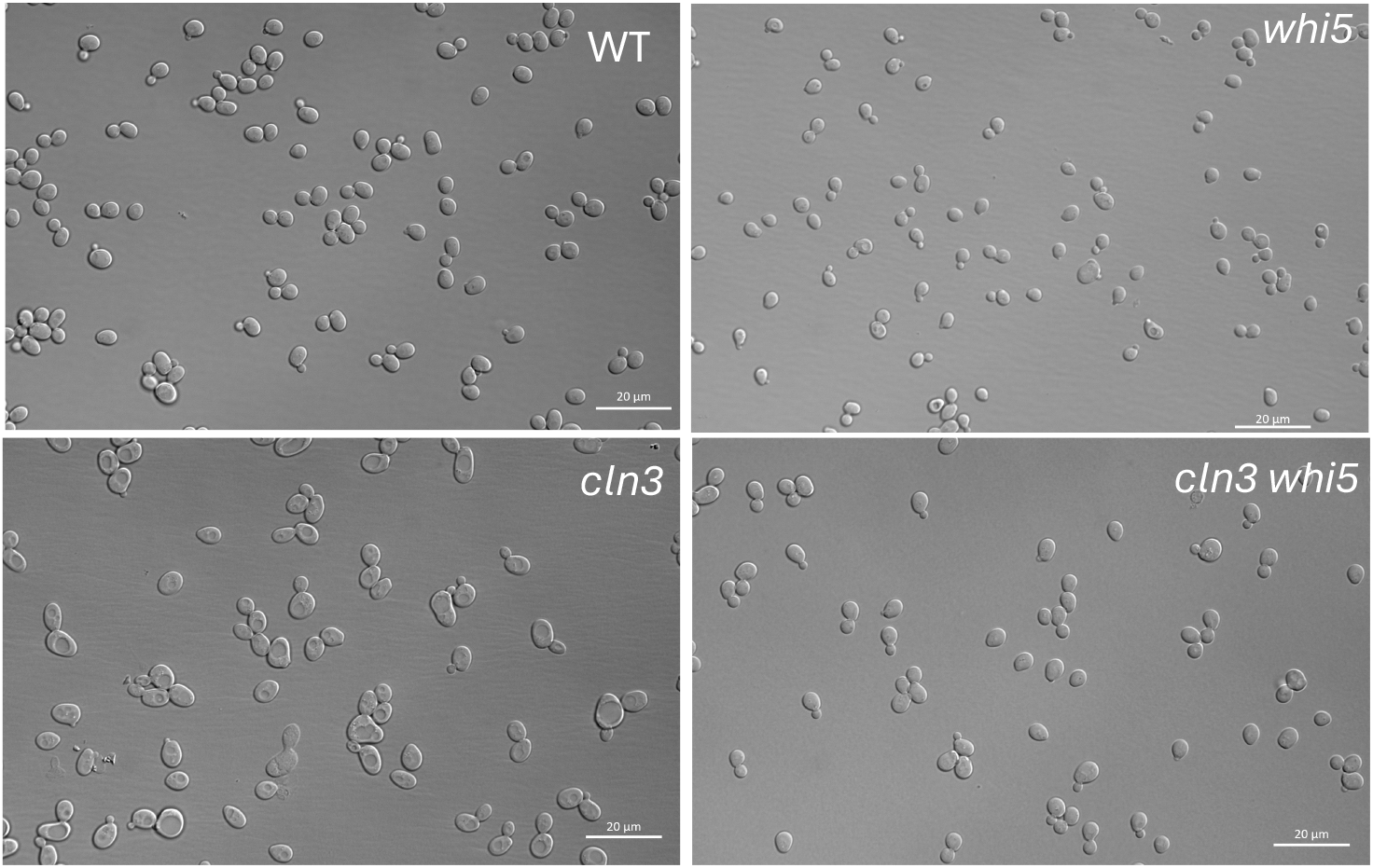
Cells in the BY background were grown to early log phase (∼2 x 10^7 cells/ml) in YEPD, and spread on agar pads on microscope slides and photographed using Nomarski optics.

Close inspection of the cell size distributions shows that the *whi5* mutant has slightly more very large cells than the wild-type, despite having smaller mean, median, and mode cell sizes. Evidence for this is shown in **Fig. 4**, which shows the right-hand tail (the large cells) from a wild-type and a *whi5* cell size distribution. At very large sizes, there are proportionately more *whi5* cells than wild-type cells, even though the mean size of the *whi5* cells is smaller. This “crossover” (more large *whi5* cells than WT cells) is not seen in all *whi5* vs WT cell culture comparisons, but it is seen in many of them. This suggests that the *whi5* mutant, at least, has some special size-control defect leading to large cells in a small proportion of cells. These might be failed attempts at “Start”, for instance. We enriched for the very large cells in a *whi5* and in a control wild-type population using a sorting flow cytometer, and tested viability. Viabilities were very high, with no significant difference in viability between the two genotypes.

**Figure 4.**
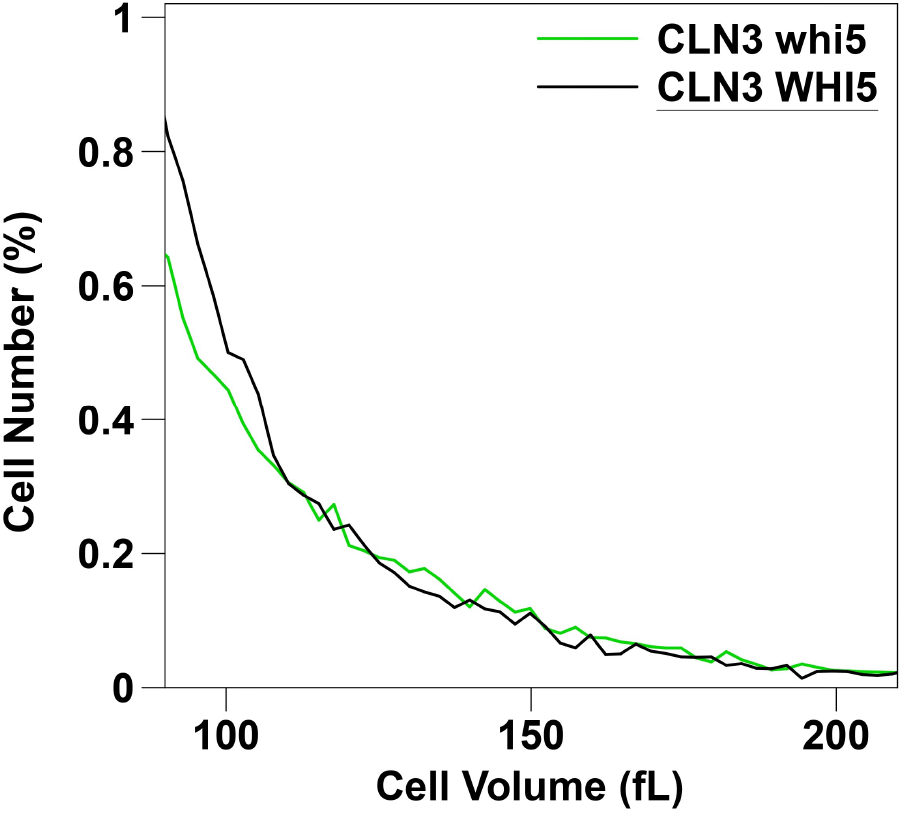
WT and *whi5* cells were grown to early log phase (∼2 x 10^7 cells/ml) in YEPD and sizes were measured using a Beckman-Coulter Multisizer. The portion of the size distribution with cell sizes between 90 fL and 210 fL is shown.

The *cln3* and *cln3 whi5* mutants also have more very large cells than wild-type, but this is expected because they also have a larger mean cell size. Therefore, we compared the four mutants for the proportion of cells that are larger than the (mean cell size + 2 standard deviations), using the mean and standard deviation for each genotype (**Table 3**). This comparison shows some evidence that the mutants have a small excess of relatively very large cells. However this difference is difficult to interpret. The size of the effect is small, and statistical significance is uncertain because the cell size distributions are not normal, so it is unclear if the criterion of “mean cell size + 2 standard deviations” is appropriate.

### Growth rate and cell size

It is not clear why yeast cells (or any other cells) are exactly the size they are. Perhaps size is selected and optimized for growth rate and doubling time. We addressed this issue with these isogenic strains of different cell sizes. Measurements included 8 independent spore clones of each genotype, except for *cln3 WHI5*, where there were 7 independent spore clones (31 spore clones total). Three independent growth-rate experiments were done, of which the most detailed was the third experiment, which is shown in Table 4. Experiments 1 and 2 are shown in detail in supplementary results, and the three experiments are summarized below in Table 5. In all three experiments the wild-type *CLN3 WHI5* strain grew fastest, while the *cln3 WHI5* mutant grew slowest. The *CLN3 whi5* and *cln3 whi5* strains were similar (and not statistically different) in growth rate; in experiment 1, the *CLN3 whi5* strain was slightly faster, while in experiments 2 and 3 it was slightly slower.

**Table 4.** Doubling Times, Expt. 3. Strains were isogenic, in the GZ background. The tetrads used were AS9, AS10, and AS12 (Table S1). Dilutions were done into pre-warmed medium. For the diploid, there were four technical replicates, which used the heterozygous parental diploid.

| Genotype | Size | Double Time | Bio. Reps. | Tech. Reps | SD | Pval: WT vs | Pval: whi5 vs | Pval: cln3 vs |
| --- | --- | --- | --- | --- | --- | --- | --- | --- |
| *CLN3 WHI5* | 49.7 | 76.1 | 3 | 2 | 0.72 |  |  |  |
| *CLN3 whi5* | 46.0 | 77.6 | 3 | 2 | 0.34 | 1e-3 |  |  |
| *cln3 WHI5* | 75.6 | 82.6 | 3 | 2 | 0.53 | 7e-9 | 3e-9 |  |
| *cln3 whi5* | 56.6 | 77.4 | 3 | 2 | 0.60 | 7e-3 | ns | 2e-8 |
| Diploid | 89.2 | 73.6 | 1 | 4 | 0.44 |  |  |  |

**Table 5.** Summary of three doubling time experiments. Three independent doubling time experiments were done, by BF (E1) and AS (E2, E3). Doubling Times are in minutes. “N” gives the total number of independent yeast cultures measured for each genotype. P-values were calculated separately for each experiment, then combined using Fisher’s method. “ns” is “not significant”. All strains were in the GZ background. E1 had several technical differences from E2 and E3 (see Supplementary data), including the use of pre-warmed medium in E2 and E3, but not E1.

| Genotype | Doubling Times |  |  | N | Pval vs WT | Pval vs whi5 | Pval vs cln3 |
| --- | --- | --- | --- | --- | --- | --- | --- |
|  | E1 | E2 | E3 |  |  |  |  |
| CLN3 WHI5 | 93.6 | 73.2 | 76.1 | 14 |  |  |  |
| CLN3 **whi5** | 95.8 | 74.4 | 77.6 | 14 | < 0.00001 |  |  |
| **cln3** WHI5 | 105.5 | 79.1 | 82.6 | 13 | < 0.00001 | <0.00001 |  |
| **cln3 whi5** | 96.5 | 73.5 | 77.4 | 14 | < 0.0005 | ns | <0.00001 |
| diploid | 91.6 | 70.6 | 73.6 | 7 |  |  |  |

As cells grow, volume and mass increase with the cube of cell radius, while surface area only increases with the square. Therefore the surface-area-to-volume ratio decreases as cells grow, and at some point this ratio would become limiting for growth. However this ratio is not necessarily limiting for growth at small cell sizes. Since nutrients typically enter cells via dedicated cell-surface transporters, and since large cells might have a higher density of transporters on their membranes, there might not be any effect of the surface-to-volume ratio on cell growth rate, as long as cells remain sufficiently small. Indeed, haploid and diploid yeast have about the same maximum growth rate, despite the fact that diploids are about twice the size of haploids (e.g., (Mable 2001; Harari, Ram and Kupiec 2018)).

Consistent with this previous work, Tables 4 and 5 also suggests that the surface-to-volume ratio is not limiting. The largest cells (i.e., the cells with the smallest surface-to-volume ratio) are the diploids, and yet these have the shortest doubling time. The sharp decrease in doubling time from the haploid *cln3* mutants (size, 75.6 fL, doubling time, 82.6 min) to the diploid strains (size, 89.2 fL, doubling time, 73.6 min) is more consistent with the idea that the cell-volume-to-genome ratio is limiting at these sizes ((Zhurinsky *et al*. 2010; Neurohr *et al*. 2019; Mu *et al*. 2020; Lanz *et al*. 2022; Swaffer *et al*. 2023; Lessenger *et al*. 2025)). At these sizes, doubling the genome allows faster growth. The idea that the volume-to-genome ratio is limiting is consistent with the sizes and growth rates of the diploid (size 89.2, doubling time 73.6, two genomes), the wild-type (size 49.7, doubling time 76.1), the *cln3 whi5* mutant (size 56.6, doubling time 77.4), and the *cln3* mutant (size 75.6, doubling time 82.6). That is, in this set of strains, the strain with the best (smallest) volume-to-genome ratio, the diploid, grows fastest, and the strain with the second best ratio, the wild-type haploid, grows second fastest, and so on.

However, the *whi5* mutant requires a different explanation. It has the smallest cells, and yet doubles slightly more slowly than wild-type. This was true in all three growth rate experiments, and in fact was also observed by Jorgensen et al. (Jorgensen *et al*. 2002), who first isolated *whi5*. Presumably at some small size, cells become “too small” in some sense. K. Dill (personal communication) has suggested that very small cells cannot help but contain a too small number of copies of some rare, important proteins, and that this leads to cellular noise. No doubt there are other possibilities. Thus, overall, these observations are consistent with the idea that wild-type cells have the optimum size for fastest growth, as one might expect from evolution. Larger cells might be limited for genome; smaller cells might be limited for something else, perhaps the copy number of rare proteins.

Another possibility is that the *cln3* and *whi5* mutations are affecting growth rate independently of cell size. That is, it is possible that a *cln3* mutant might grow slowly because of a lack of G1 CDK kinase activity, independently of cell size. Similarly, a *whi5* mutant might grow slowly because of a lack of Whi5 activity, independently of cell size. But in fact, the slow doubling time of the *cln3* mutant is suppressed by the *whi5* mutation (i.e., the *cln3 whi5* double mutant grows faster than the *cln3*), and the slow doubling of the *whi5* mutant is at least not made worse by adding the *cln3* mutant. These results argue the slow growth of the *cln3* mutant, and possibly the *whi5* mutant, is indeed due to its cell size, and not directly due to a lack of Cln3 or Whi5 activity.

## Discussion

We have three major findings. First, contrary to some simplified cell cycle models, neither *CLN3* nor *WHI5* is epistatic to the other. However, they are also probably not completely independent. A likely interpretation is that *CLN3* does work in part via *WHI5*, but also has very significant *WHI5*-independent mechanisms. These *WHI5*-independent pathways for the Cln3-Cdc28 kinase might include phosphorylation of the C-terminal domain of RNA polymerase II (Koivomagi *et al*. 2021), or the phosphorylation of Stb1 (Wang *et al*. 2009).

Second, the individual phenotypes of *cln3* and of *whi5* largely cancel each other in the double mutant. The *cln3 whi5* double mutant is almost wild-type in size, morphology, growth rate, and cell size coefficient of variation. Cells regulate size and division very well even in the complete absence of both of these two major cell size regulators. This is perhaps our most surprising finding: what is the point of these two major cell cycle regulators, if the cell is nearly wild-type when lacking both of them? Of course, there may be other significant phenotypes in the double mutant that we have not noticed. Still, the almost wild-type doubling time suggests other phenotypes cannot be very severe, at least under standard YEPD growth conditions.

Another cell cycle activator in yeast is *BCK2*, and phenotypes of the *bck2* mutant have some similarities to those of the *cln3* mutant. Interestingly, Chadha et al. (Chadha *et al*. 2024) recently found that the *bck2 whi5* double mutant has a size quite similar to wild-type, and has surprisingly efficient size homeostasis, considering that it lacks two major cell size controllers. In these ways, the *bck2 whi5* double mutant is similar to the *cln3 whi5* double mutant.

Third, all three of the mutants, whether they are large (*cln3, cln3 whi5*) or small (*whi5*) grow more slowly than wild-type, consistent with the idea that the wild-type has optimized cell size for growth rate. The growth rate differences are small, but we have used isogenic strains; we have used multiple different, independent isolates of each genotype; we have done many technical replicates with each isolate; and we have sampled some different growth conditions. Consistently, and with very small p-values, all three mutants grow more slowly than the wild-type. The slow growth of large cells is consistent with the genome being limiting at large cell sizes, particularly since the diploid, with an extra genome, grows faster than the wild-type haploid, even while being 79% bigger (89.2/49.8, Fig. 2). However, the slow growth of the small *whi5* cells requires some different explanation. While there are many possible reasons why a cell might be “too small”, one attractive and testable possibility is that a small cell will have too few copies of important, rare proteins, and so will suffer from noise in the face of stochastic variation of protein copy number (Dill, pers. comm.).

While it is in some ways obvious that a diploid cell should be about twice as big as a haploid cell, since it has twice as much of everything, still, the mechanism by which diploids become twice as big as haploids is unclear. But the reasoning above at least provides a rationale for the diploid-haploid size difference. There is some penalty for being “too small” (perhaps noise from variation in copy number), and also a penalty for being “too big”, which is likely a large cell-size-to-genome ratio. Diploids have an extra genome; therefore almost doubling in size allows them to minimize the “too small” penalty, without increasing the “too big” penalty.

If one imagines a strain of yeast over evolutionary time, it is presumably evolving to maintain the optimum cell size. And yet, depending on other genes in the genome, and on environmental conditions and available nutrients, etc. etc., that optimum size is likely to vary. And so, as time passes, there will be a succession of mutations that fine-tune cell size to the current optimum. There will be an accretion of cell size control alleles. And so, when one looks at the function of particular cell size control alleles, one may be looking into the evolutionary past, when the optimum size went up, or down, or up and then down.

## Acknowledgements

We thank Janet Leatherwood for advice throughout this work, and for reading the manuscript. We thank Ken Dill for suggesting the protein copy number problem as a problem limiting cell size on the small side, Kurt Schmoller for telling us the phenotype of the *bck2 whi5* double mutant prior to publication, and Kara Swenson for help with microscopy. This work was funded by R01 GM127542 to BF.

## Supplementary Tables S1 and S2

**Table S1.** Cell sizes in the GZ background, in fL.

| **Background: GZ238/GZ239** |  |  |  |  |  |  |
| --- | --- | --- | --- | --- | --- | --- |
| N | Genotype | Mean (SE) | Median (SE) | Mode (SE) | C.V. | Strain |
| 6 | *CLN3 WHI5* | 50.6 (0.06) | 45.3 (0.00) | 35.1 (0.89) | 0.46 | AS-9D |
| 6 | *CLN3 whi5* | 46.6 (0.34) | 40.6 (0.25) | 29.7 (0.51) | 0.54 | AS-9C |
| 6 | *cln3 WHI5* | 76.1 (0.32) | 69.5 (0.32) | 54.7 (1.41) | 0.44 | AS-9A |
| 6 | *cln3 whi5* | 57.8 (0.18) | 51.2 (0.00) | 38.3 (0.91) | 0.48 | AS-9B |
| 6 | *CLN3 WHI5* | 49.5 (0.26) | 44.8 (0.32) | 34.4 (0.91) | 0.46 | AS-10A |
| 6 | *CLN3 whi5* | 45.5 (0.20) | 39.3 (0.00) | 28.7 (0.44) | 0.54 | AS-10D |
| 6 | *cln3 WHI5* | 76.4 (0.33) | 69.8 (0.34) | 58.6 (1.59) | 0.44 | AS-10C |
| 6 | *cln3 whi5* | 56.6 (0.63) | 50.5 (0.51) | 38.8 (0.73) | 0.48 | AS-10B |
| 6 | *CLN3 WHI5* | 49.2 (0.32) | 43.8 (0.00) | 34.1 (0.99) | 0.49 | AS-12B |
| 6 | *CLN3 whi5* | 45.8 (0.18) | 39.6 (0.25) | 30.2 (0.80) | 0.53 | AS-12A |
| 6 | *cln3 WHI5* | 74.4 (0.41) | 67.8 (0.46) | 52.4 (2.64) | 0.44 | AS-12C |
| 6 | *cln3 whi5* | 55.4 (0.21) | 48.2 (0.00) | 37.4 (1.05) | 0.50 | AS-12D |
| 8 | Diploid | 89.2 (0.38) | 83.6 (0.33) | 66.6 (0.96) | 0.37 | Diploid |

**Table S2.** Cell sizes in the BY background, in fL.

| **Background: BY4741/BY4742** |  |  |  |  |  |  |
| --- | --- | --- | --- | --- | --- | --- |
| N | Genotype | Mean | Median | Mode | C.V. | Strain |
| 1 | *CLN3 WHI5* | 38.4 | 35.7 | 29.9 | 0.39 | YSH 675-2A |
| 1 | *CLN3 **whi5*** | 30.4 | 27.8 | 24.2 | 0.41 | YSH 675-2D |
| 1 | ***cln3** WHI5* | 61.2 | 55.9 | 43.0 | 0.42 | YSH 675-2C |
| 1 | ***cln3 whi5*** | 43.4 | 40.0 | 32.8 | 0.41 | YSH 675-2B |
| 1 | *CLN3 WHI5* | 38.1 | 35.7 | 31.4 | 0.39 | YSH 675-5C |
| 1 | *CLN3 **whi5*** | 33.4 | 30.7 | 24.9 | 0.43 | YSH 675-5D |
| 1 | ***cln3** WHI5* | 64.2 | 58.8 | 49.4 | 0.44 | YSH 675-5A |
| 1 | ***cln3 whi5*** | 40.6 | 37.2 | 30.7 | 0.44 | YSH 675-5B |
| 1 | *CLN3 WHI5* | 37.2 | 35.0 | 29.9 | 0.38 | YSH 675-6A |
| 1 | *CLN3 **whi5*** | 30.4 | 28.5 | 25.6 | 0.39 | YSH 675-6B |
| 1 | ***cln3** WHI5* | 65.0 | 59.5 | 46.5 | 0.44 | YSH 675-6D |
| 1 | ***cln3 whi5*** | 41.4 | 38.6 | 31.4 | 0.40 | YSH 675-6C |

